# Optical Screening Identifies Chemical Modulators of Intracellular α-synuclein Aggregation

**DOI:** 10.64898/2026.07.04.736150

**Authors:** Lily Rothschild, Colter Giem, Aditya Bajaj, Jessica W. Luo, Kimberly L. Carey, Jacques Deguine, Ramnik J. Xavier

## Abstract

Parkinson’s disease (PD) is a movement disorder characterized by the accumulation of alpha-synuclein aggregates leading to dopaminergic neuron loss in the substantia nigra. While PD has been associated with environmental and microbiome changes, our ability to assess the mechanistic impact of these factors on synuclein aggregation in cells has remained limited. Here, we designed and optimized a high-throughput optical screening system to assess the effect of metabolites and small molecules on synuclein aggregation in cell lines expressing a synuclein-fluorescent protein fusion and treated with pre-formed fibrils (PFFs). Using this assay, we identified several compounds that modulate synuclein aggregate accumulation in cells, including harman, a β-carboline that led to reduced synuclein aggregation. We further investigated the transcriptional effect of harman and PFFs and identified changes in peroxiredoxins as a potential mechanism linking harman to aggregate accumulation. Altogether, this work establishes a pipeline to prioritize small molecules that can impact synuclein aggregate formation.

## Introduction

Parkinson’s disease (PD) is a neurodegenerative disorder most distinctly characterized by the loss of dopaminergic neurons in the substantia nigra^1^. Many epidemiological and genetic association studies have highlighted the complex etiology of PD, which includes familial and sporadic cases that can be driven by mutations in alpha-synuclein (α-syn, encoded by *SNCA*) ^2^, *LRRK2*, *PRKN* ^3^, *PINK1* and others^4^; or, in the more common sporadic cases, environmental toxicants ^5,6^.

Regardless of etiology, the majority of PD cases show an expansion of proteinaceous inclusion bodies, including α-syn aggregates. Together with human genetic data, animal models based on the over-expression of mutated synucleins (A53T, A30P) ^7^ all suggest a causal role for α-syn mutation and inclusion body formation in neurodegeneration and PD pathology. While PD symptoms appear to be primarily driven by the loss of dopaminergic neurons, α-syn aggregates have been detected in many other regions both in and out of the brain, and a recent study showed that testing for phosphorylated synuclein in skin biopsies can identify PD and related disorders with high specificity and sensitivity ^8^. The identification of synuclein aggregates in enteric neurons and the dorsal motor of the vagus led to the development of the Braak hypothesis ^9^, proposing a “body-first” subtype of the disease with early peripheral nervous system involvement ^10,11^. Consistent with this hypothesis, functional gastrointestinal disorders often precede the development of motor symptoms in PD ^12,13^. An animal model based on gastrointestinal injection of pre-formed synuclein fibrils (PFF) demonstrated that pathological aggregates can spread to the brain through the vagus nerve ^14,15^, lending support to this mechanism as one possible route of PD development.

Large scale studies of PD patient cohorts have highlighted numerous changes in the intestinal microbiome of these patients, including in the earliest phases of disease ^16,17^. While these shifts are difficult to disentangle from those associated with constipation, murine models of disease have suggested that intestinal microbes can directly contribute to synucleinopathy progression ^18,19^, including potentially through microbe-derived amyloids. The microbiota also interacts with the host through a complex collection of small molecules and metabolites ^20^ that we and others have begun to characterize through a combination of metagenomic and metabolomic approaches ^21–23^. In PD in particular, studies have pointed out decreases in short chain fatty acid abundance, with potential effects on intestinal transit and inflammation ^24,25^, although the extent of differential metabolites and their impact on disease onset and progression remains largely uncharted.

Here, we set out to assess the impact of microbiota-associated metabolites on synuclein aggregation by developing a high-throughput screening platform based on GFP-α-syn expressing cells and a test library of metabolites composed of molecules identified through the Human Microbiome Project (HMP). We identify a number of molecules associated with increased or decreased synuclein aggregation in this assay and further characterize the effects of an inhibitor of synuclein aggregation through transcriptomic profiling. Altogether, our findings establish a robust platform to detect and quantify α-syn aggregation and establish that gut derived metabolites can modulate this process, opening new avenues to mechanistically characterize microbes associated with PD.

## Results

### Establishment of an α-syn-GFP HEK293T Cell Line for Quantitative Analysis of Aggregation

We first set out to establish a high-throughput assay for the detection of synuclein aggregation. Prior studies demonstrated that a GFP-a-syn fusion expression construct does not spontaneously self-aggregate ^26^ but can enable the detection of induced aggregation as regions of high GFP intensity ^27^. We therefore first established stable HEK293T cells expressing this fusion protein, sorted transduced cells and selected single cell clones, confirming uniform and robust GFP expression in the α-synuclein-GFP HEK293T (α-syn-GFP) selected cell line compared to controls (**Fig. 1A**). Next, we aimed to validate the ability of these cells to report induced synuclein aggregation by transfecting them with pre-formed synuclein fibrils (PFFs) for 24h. Cells were subsequently fixed and imaged on a high-throughput confocal microscope. As expected, treatment with PFFs induced the formation of discrete punctate GFP spots, while untreated cells retained diffuse fluorescence (**Fig. 1B**).

**Figure 1.**
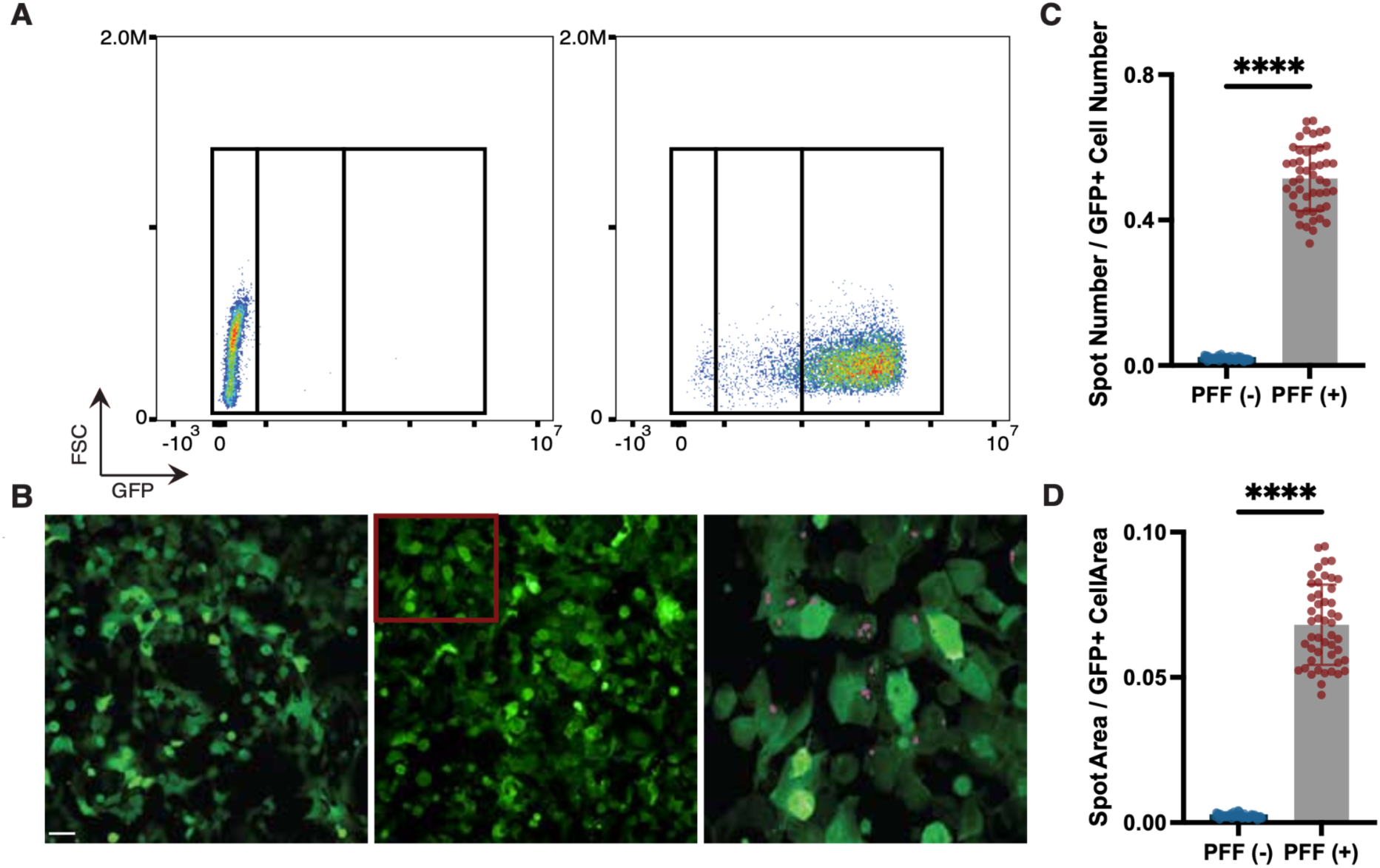
An α-synuclein-fluorescent protein fusion construct provides an aggregation screening platform. **(A)** Flow cytometry analysis used to generate a stable α-syn-GFP HEK293T cell line. Shown are representative FACS plots from GFP-negative controls HEK293T cells (left) and α-syn-GFP positive cells (right). **(B)** Representative confocal images of α-syn-GFP cells untreated (left) or treated with preformed fibrils (PFFs; middle), and a magnified view highlighting intracellular α-synuclein puncta (outlined in purple; right). Scale bar, 50 *μ*m. **(C-D)** Quantification of α-synuclein aggregation based on puncta per cell **(C)** and puncta area normalized to cell area **(D)** Unpaired two-tailed t-test with Welch’s correction: ****p < 0.0001; n = 48 wells per group; pooled from 4 independent experiments.

Using these conditions as our positive and negative controls, respectively, we developed a custom image analysis pipeline to enable the automated detection of GFP spots within cells passing filtering criteria (see **Methods**). This analysis revealed a significant increase in both puncta number per cell and puncta area normalized to total cell area following PFF exposure (**Fig. 1C-D**). To validate that the induction of fluorescent punctae was specific to the synuclein moiety of the construct, we created a second HEK293T cell line, α-syn-GFP-mCherry, that included mCherry as an untagged fluorescent protein, serving as negative control to test for nonspecific punctae formation (**Fig. S1A**). We confirmed uniform and robust GFP and mCherry expression in the α-syn-GFP-mCherry HEK293T cell line compared to control HEK293T cells and α-syn-GFP HEK293T cells by flow cytometry (**Fig. S1B**) and treated and imaged the cells with PFF as described above (**Fig. S1C**). We then separately applied our analysis to the GFP and mCherry channels and confirmed the specificity of punctae formation after PFF treatment to the GFP channel, where mCherry signals showed little to no punctae formation in either condition (**Fig. S1D-E**). Finally, to confirm the suitability of this assay for screening, we determined a Z-prime score for this assay based on the measurement of either spot number per cell or spot area per cell area, yielding Z’ = 0.433 and Z’ = 0.337, respectively, suggesting that the assay is generally applicable for screening even if it suffers from some level of noise ^28^. Altogether, these results establish the α-syn-GFP HEK293T line as a robust and scalable model for detecting and quantifying intracellular α-syn aggregation, which can be applied to high throughput screening.

### High-Throughput Screening in HEK293T Cells Identifies Candidate Modulators of α-synuclein Aggregation

To identify intestine associated compounds that modulate intracellular α-syn aggregation, we screened a library of 318 individual compounds, selected from metabolomic data generated from human stool and controls with known bioactivity, at stock concentrations of 1 or 10 mM in DMSO, depending on solubility (**Supplementary Table 2**). GFP-α-syn HEK293T were treated with compounds at a 125-fold dilution (corresponding to 0.8% DMSO final) then with PFFs in replicate plates. Cells were fixed and imaged after 24h and the resulting data was analyzed as described above to determine the number and area of spots induced by PFFs in the different treatment conditions. We assessed the impact of compounds on the formation of spots relative to DMSO, and identified multiple compounds that reproducibly increased aggregation, measured either by spot number (**Fig. 2A**) or area (**Fig. 2B**), alongside a small set of compounds that diminished spot formation. Impact of the compound on total cell numbers was relatively limited (**Fig. 2C**), suggesting that toxicity was not a major driver of changes in spot detection.

**Figure 2.**
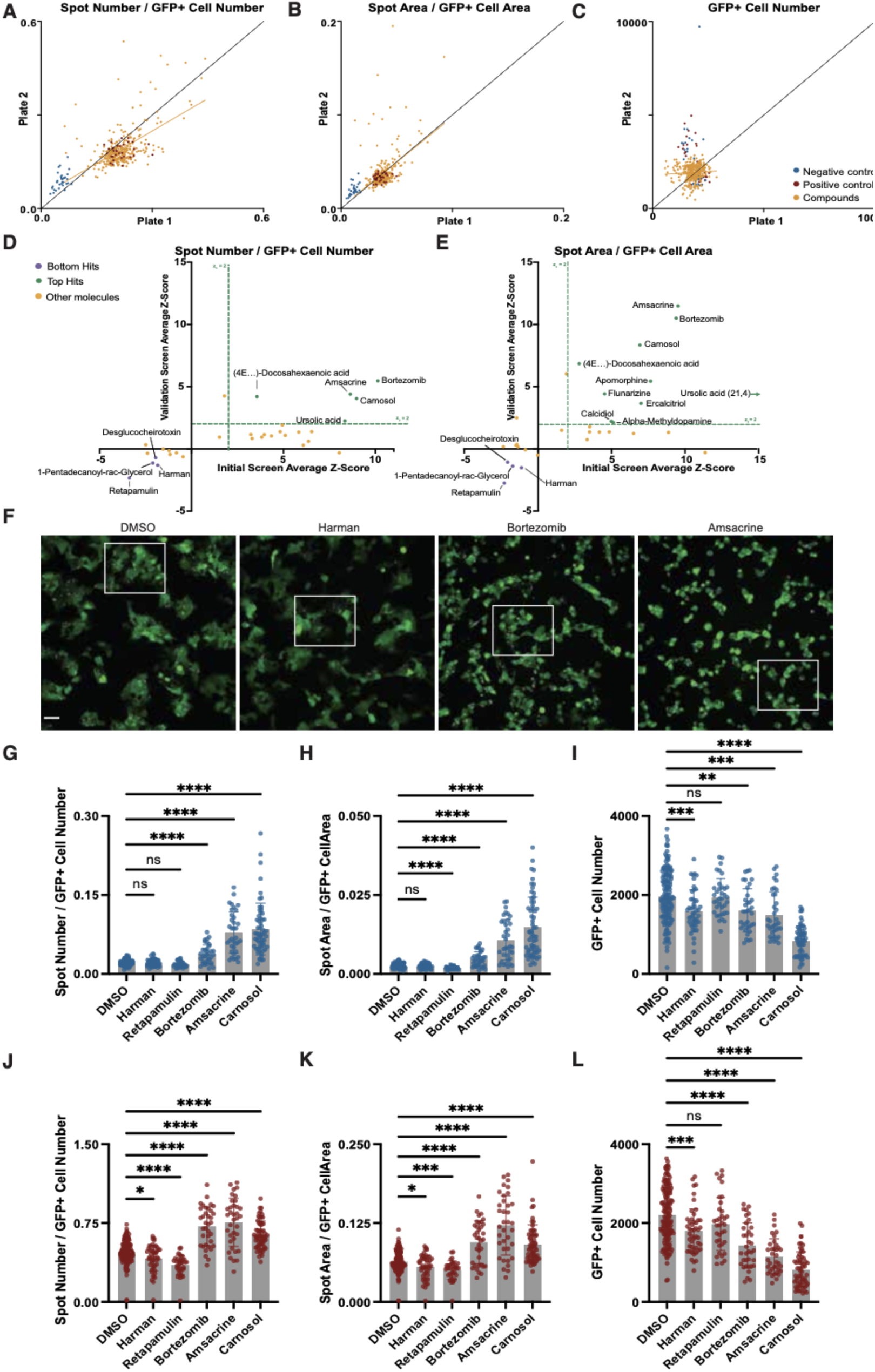
High-throughput screen of intestine-associated compounds identifies modulators of α-synuclein aggregation (A-C) Initial screen results for 320 microbiome and gut-related compounds applied at concentrations of 8 or 80 μM across replicate plates in one experiment representative of two individual screens; intracellular GFP puncta are quantified as GFP spot number normalized to cell number **(A)**, GFP spot area normalized to cell area **(B)**, and GFP+ cell number **(C)**. Positive controls (red), negative controls (blue), and compounds (yellow) are annotated in the legend. **(D-E)** Triplicate validation of 32 screen hits assessed across replicate plates by GFP puncta number per cell **(D)** and GFP puncta area per cell area **(E)** relative to the initial screen z-scores. Inducers with a z-score ≥ 2 are labeled in orange and reducers with a z-score ≤ 0.5 are labeled in purple. **(F-L)** Representative confocal microscopy images. White boxes indicate regions shown at higher magnification in corresponding zoomed panels (**Fig. S2E**) **(F)** and quantification of GFP puncta number per cell **(G,J)**, GFP puncta area per cell area **(H,K)**, and GFP cell number **(I,L)** of α-syn-GFP HEK293T cells in the absence of PFFs (**G-I**; blue) or in the presence of PFFs (**J-L;** red), with the addition of DMSO (n = 144 wells), Harman (n = 48 wells), Retapamulin (n = 36 wells), Bortezomib (n = 36 wells), Amsacrine (n = 36 wells), or Carnosol (n = 60 wells); (pooled from 3 independent experiments with 2 replicate plates each). Scale bar, 50 μm. *p < 0.05, **p < 0.01, ***p < 0.001 ****p < 0.0001. Data analyzed by ANOVA with Dunnett’s T3 multiple comparisons post-hoc test.

To quantitatively determine the impact of candidate modulators, we calculated z-scores for each compound within individual plates, by comparing the value of each compound to the distribution of DMSO-treated wells. We identified a set of aggregation promoting compounds with z-scores ≥ 2 across metrics. Our ability to identify compounds that reduce aggregation was partially limited by the dynamic range of the assay. To assess this, we first calculated z-scores for spot formation in cells without PFFs, which would correspond to no induction of synuclein aggregation. We observed a range of -6.08 to -2.78 for spot number normalized to GFP+ cell number and -5.74 to -1.49 for spot area normalized to GFP+ cell area, suggesting that a z-score at or below -1.5 would represent a complete abrogation of aggregation. Therefore, we included all compounds with z-scores ≤ -0.5 as candidate inhibitors given the more limited dynamic range of the assay in this direction. Hits were defined as compounds that met these criteria in both puncta number per cell and puncta area normalized to cell area across replicate plates in two independent experiments. Using this approach, we identified 32 primary hits that reproducibly increased or decreased α-syn aggregation (**Supplementary Table 2**). To validate these initial findings, the 32 compounds were retested in triplicate in replicate plates.This experiment validated reproducible effects for five aggregation promoting compounds – amsacrine, bortezomib, carnosol, ursolic acid, and (4E,7E,10E,13E,16E,19E)-4,7,10,13,16,19-Docosahexaenoic acid – and four aggregation reducing compounds – Desglucoheirotoxin, Harman, Retapamulin (all at 80 μm), and 1-Pentadecanoyl-rac Glycerol (at 8 μM), (**Fig. 2D-E, Fig. S2A-B**). We also tested these compounds at a 5-fold dilution and only bortezomib, a known proteasome inhibitor, still showed a significant effect at 16 μM, an observation consistent with its impact on proteasome activity at concentrations below 1μM across cell lines ^29^ (**Fig. S2C-D**).

Select top and bottom hits with the most significant effects on synuclein aggregation were used for testing in a larger replication set. We examined and validated the effect of harman – a heterocyclic amine and ꞵ-carboline found in multiple foods and in tobacco smoke – and retapamulin – a topical antibiotic derived from fungi – as top reducers, whereas carnosol – a dietary terpene from rosemary ^30^ – bortezomib, and amsacrine were identified as top inducers across the replication set (**Fig. 2F-L**).

Cells treated with harman displayed markedly fewer and smaller puncta, and two representative aggregation promoting metabolites, bortezomib and amsacrine, displayed increased and larger puncta compared with the discrete aggregates observed in DMSO-treated PFF positive controls (**Fig. 2F-H, 2J-K, Fig. S2E**). We also evaluated changes in cell numbers across these experiments and observed that treatment with most compounds had modest but significant effects on cell counts, which were largely similar in the absence and presence of PFFs (**Fig, 2I, 2L**). Overall, compounds associated with increased aggregation showed a higher reduction in cell counts, although this was also true in the absence of PFFs. Conversely, bortezomib and harman, which have opposite effects on aggregation, did not alter counts in the absence of PFFs, but the former showed a slight increase in cell number reduction in the presence of PFFs (q = 0.026), potentially tying this reduction to aggregate formation.

Altogether, our screening approach identified several compounds linked to increased synuclein aggregation, and our measurements of cell viability suggests increased cell stress as a potential mechanism. Interestingly, we also identified a small number of compounds linked to reduced spot formation, suggesting that these could be associated with decreased aggregation or increased clearance.

### Transcriptomic Profiling of Harman and PFF Treated Cells

To understand the potential mechanisms behind this reduction in spot formation, we focused on harman and performed bulk RNA-seq on control HEK293T cells (without synuclein expression) and α-syn-GFP-mCherry cells treated with harman, PFFs, or both. We validated the effect of harman on synuclein aggregation in these assays (**Fig. 3A-C, Fig. S3A-C**) and analyzed gene expression 24h after treatment. Since α-syn-GFP-mCherry expresses its own synuclein and the control HEK293T cells do not, we analyzed the two lines separately. Gene expression was modeled as a function of harman treatment, fibril exposure, and the interaction between the two, accounting for batch effects with a replicate term (**Fig. 3D**).

**Figure 3.**
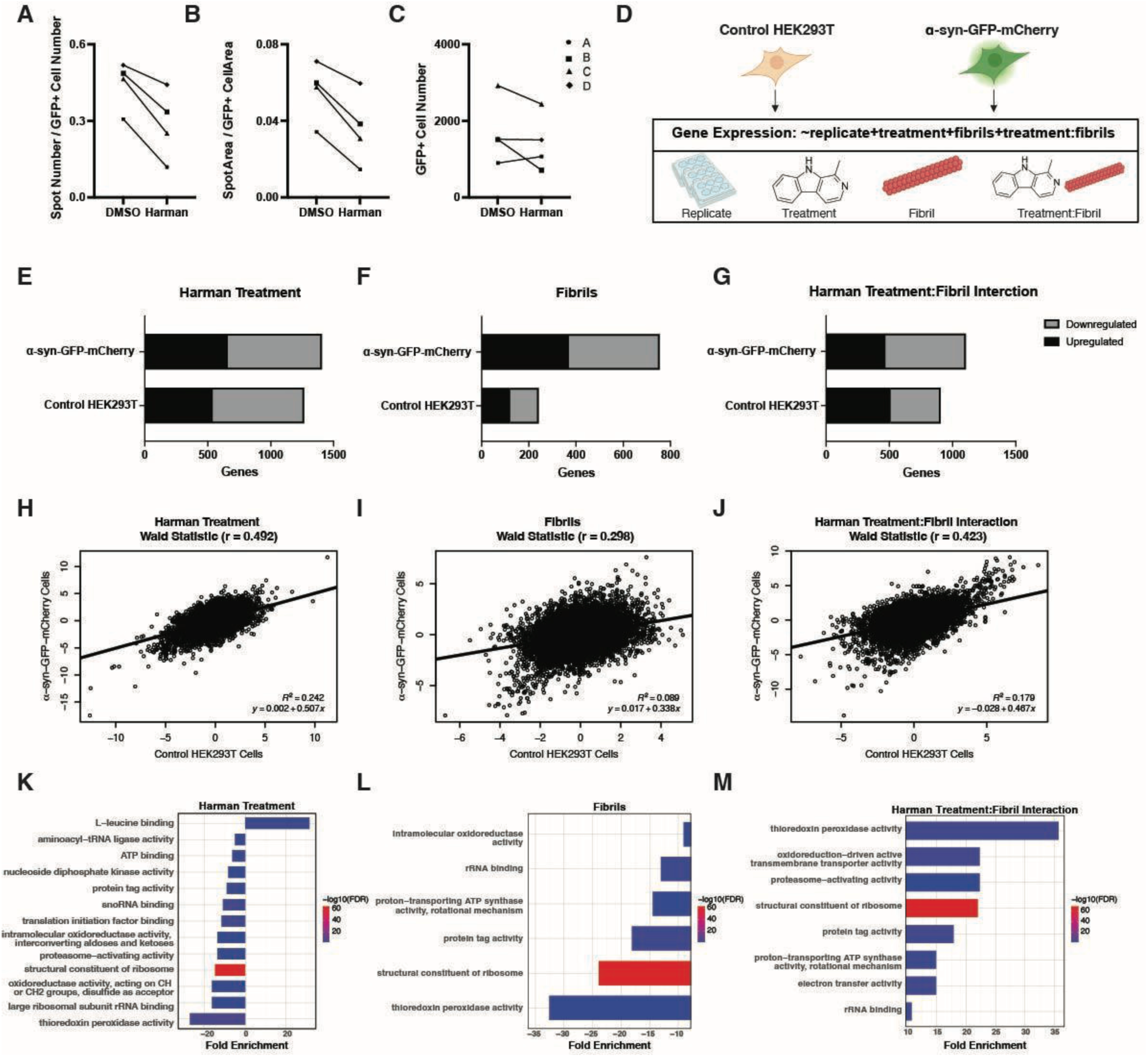
Differential expression and pathway enrichment analyses reveal transcriptional changes associated with harman exposure (A-C) Quantification of GFP puncta number per cell **(A)**, GFP puncta area per cell area **(B)**, and GFP cell number **(C)** in cells treated with PFFs and DMSO or PFFs and harman. Each point presents the mean of all wells from one independent experiment (A: DMSO = 12 wells, harman = 12 wells; C: DMSO = 24 wells, harman = 21 wells; B and D: DMSO = 24 wells, harman = 24 wells) **(D)** Schematic of linear model used for RNA-seq differential expression analysis incorporating replicate, treatment, fibrils, and a treatment:fibril interaction term created with Biorender.com. **(E-G)** Number of significant (FDR < 0.2) up- and downregulated genes in each cell line by model terms: Treatment **(E)**, Fibrils **(F)**, and Treatment:Fibrils Interaction filtered**(G)**.**(H-J)** Scatter plots of DESeq2 Wald Statistics between HEK293T control cells α-syn-GFP-mCherry cells for all genes by (H) harman treatment, (I) fibrils, and (J) their interaction labeled with Pearson correlation coefficient (r) and fit with a linear regression **(K-M)** Functionome Gene ontology (GO) molecular function analysis of genes in the α-syn-GFP-mCherry cell line by each model term: Harman Treatment **(K)**, Fibrils **(L)**, and Treatment:Fibrils Interaction **(M)** Only significant pathways with a fold enrichment ≥ 5 are represented.

We first summarized the number of significant differentially expressed genes (DEGs, FDR < 0.2) by each condition in each cell line (**Fig. 3E-G, Supplementary Table 3**). As expected, PFF exposure had a larger impact on α-syn-GFP-mCherry cells, inducing a stronger response in the presence of endogenously expressed synuclein (**Fig. 3F**). On the other hand, harman treatment induced a similar number of DEGs in both control HEK293T and α-syn-GFP-mCherry cells (**Fig. 3E**) and the same was true for the harman treatment:fibril interaction term (**Fig. 3G**).

We then assessed gene-level correlation of the transcriptional responses between the two cell lines using the Wald statistic calculated for each term. The response to harman showed a moderate correlation (Pearson r = 0.492) (**Fig. 3H**), while fibril induced responses were less correlated (r = 0.298), consistent with a synuclein-dependent effect (**Fig. 3I**) and the interaction term showed a moderate correlation (r = 0.423) (**Fig. 3J**). These results suggest that PFF-driven transcriptional responses depend at least partially on endogenous synuclein expression, and we therefore focused our analysis on α-syn-GFP-mCherry lines.

To further investigate the specific effect of fibrils in synuclein expressing cells, we looked at genes that were significantly up- or downregulated in α-syn-GFP-mCherry cells but not in control HEK293T cells after fibril treatment. We performed gene ontology enrichment molecular function analysis with the PANTHER (PAN-GO) Human Functionome ^31^ on these gene sets and observed a downregulation of pathways involved in mitochondrial function and translation specifically in synuclein expressors (**Fig. S3D**). By contrast, no enriched pathways were identified in control cells, confirming that PFFs have a stronger effect on cells that express endogenous synuclein.

### Pathway Analysis of Harman-Responsive Genes

Next, we used the same enrichment analysis to understand the effect of harman specifically in synuclein expressing α-syn-GFP-mCherry cells. We identified multiple pathways associated with harman treatment (**Fig. K**), fibrils (**Fig. 3L**), and harman treatment:fibril interaction (**Fig. 3M**), when multiple pathways of the same hierarchy were included, focused on the most specific term in the pathway hierarchy.

Changes in thioredoxin peroxidase (TPx) activity were associated with all terms, and Interestingly, we observed a downregulation of these pathways after either PFFs or harman treatment alone, while the combination of these two perturbations did not further decrease thioredoxin peroxidase activity, which was modeled as a positive association with the interaction term (i.e. the level of TPx activity in the context of harman and PFFs is higher than predicted based on each treatment considered separately). This pattern was reflected at the gene level across individual members of the TPx family including PRDX1-6 (**Fig. 4A**). Expression values measured as transcript per million showed that both harman treatment and PFF exposure reduced expression of several PRDX genes relative to untreated cells when used independently. In cells treated with both harman and fibrils, PRDX gene expression levels were generally comparable or higher than in single-treatment conditions. Cells treated with harman also showed a decrease in pathways related to ribosomal function, cytoplasmic translation, heat-shock responses, and protein folding. These patterns suggest alteration of the translational and proteostasis machinery in harman-treated cells.

**Figure 4.**
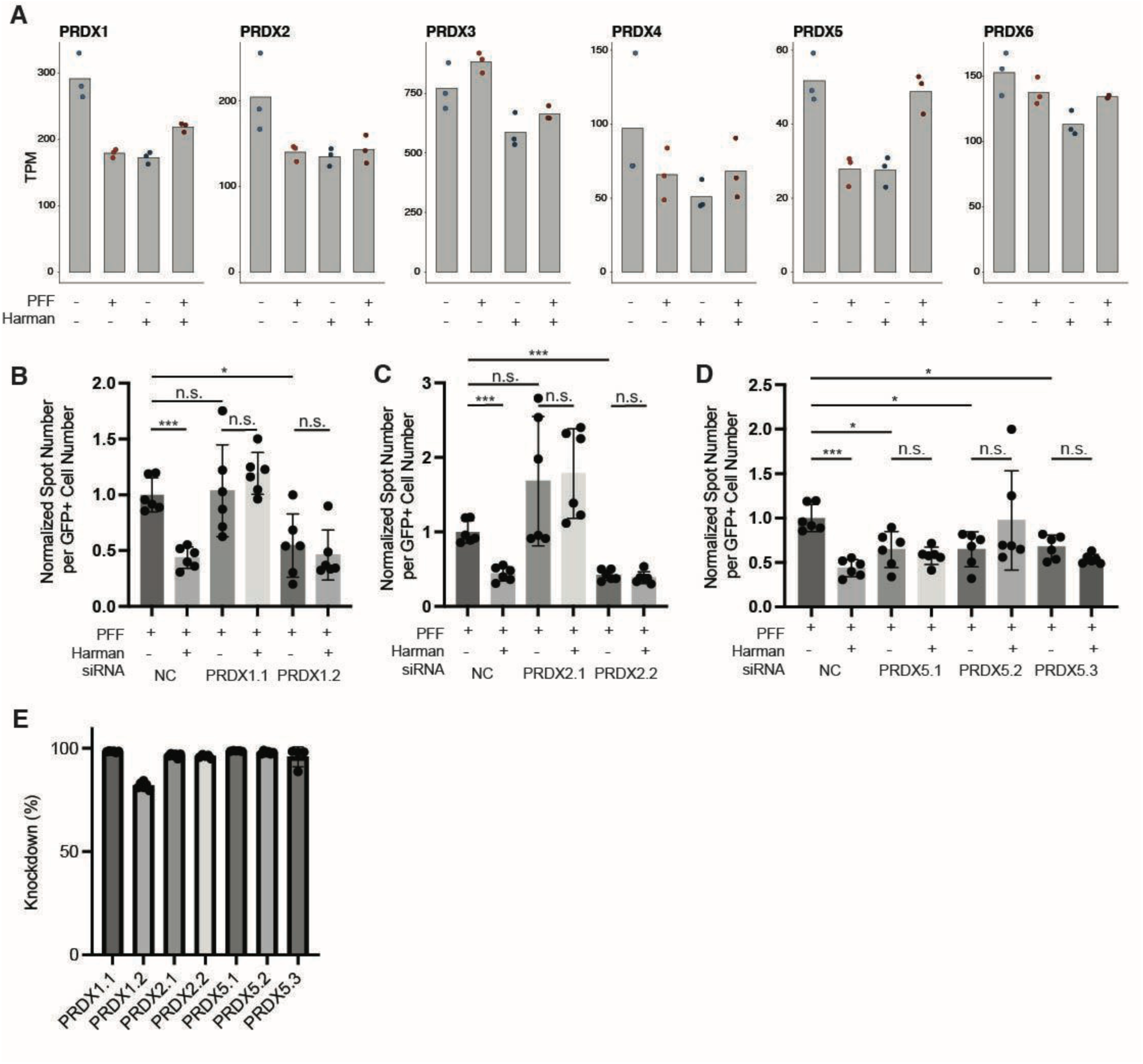
Knockdown of PRDX genes modulates synuclein aggregation. **(A)** Transcript abundance (TPM) across all isoforms of thioredoxin peroxidase activity genes (PRDX1, PRDX2, PRDX3, PRDX4, PRDX5 and PRDX6) in cells treated with or without the addition of PFFs and Harman. Values represent raw TPM counts from biological replicates. **(B-D)** Quantification of GFP puncta per GFP-positive cell cell in cells transfected with the indicated siRNA (NC = negative control or multiple PRDX1 **(B)**, PRDX2 **(C)** or PRDX5 **(D)**-targeting siRNAs), treated with PFF and with or without harman. Data is pooled from two independent experiments after normalizing to the average of the PFF only controls. Significance was assessed using a Brown-Forsythe and Welch ANOVA followed by Dunnett’s T3 multiple comparisons test (n.s. - not significant, * - p < 0.05, *** - p < 0.001) **(E)** Percentage mRNA knockdown 48h after siRNA transfection, determined by RT-qPCR compared to a non-targeting scrambled control **(H-J)** Quantification of GFP puncta per cell in cells transfected with siRNA with and without harman treatment normalized to the negative control in 2 independent experiments. PRDX1 **(H)**, PRDX2 **(I)**, PRDX5 **(J)**.

### Validation of the Impact of Thioredoxin Peroxidase Activity on Aggregation

To validate the functional role of this gene family, we used siRNAs to knock down PRDX1, PRDX2 or PRDX5 before exposing cells to PFF in the presence or absence of harman (**Fig. 4B-E**). We selected those three genes as they showed the most consistent downregulation in the presence of either harman or PFF (**Fig. 4A**). In spite of relatively similar knock-down efficiency across multiple siRNAs (**Fig. 4E**), knock-down of either PRDX1 **(Fig. 4B)** or PRDX2 **(Fig. 4C)** showed variable effects on the accumulation of synuclein aggregates within cells. By contrast, knockdown of PRDX5 with 3 independent siRNAs led to a consistent significant reduction in synuclein aggregates (**Fig. 4D**, p = 0.045, 0.042 and 0.017 for PRDX5.1, PRDX5.2 and PRDX5.3, respectively). Interestingly, harman did not further reduce synuclein aggregation in these cells, suggesting that knockdown of PRDX5 activity may be sufficient to replicate the protective effect of harman.

Together, these results suggest that harman induces a state characterized by alterations in redox and proteostatic activity which may contribute to its protective effects in the context of fibril exposure in our assay.

## Discussion

In this study, we present a high-throughput optical screening method to identify chemical modulators of intracellular α-syn aggregation in HEK293T cells expressing a GFP-α-syn fusion protein construct. This approach enables scalable, quantitative assessment of aggregation phenotypes in live cells and provides a system to interrogate how small molecules – derived from dietary or microbial sources – can influence seeded α-synuclein pathology. One important aspect of this platform is that the measured changes in aggregation can be driven by changes in either the formation of fibrils themselves, or in the clearance of aggregates, while in vitro, cell-free aggregation assays focus solely on the former ^32^. This is evidenced by the impact of bortezomib, a proteasome inhibitor, in our screen, as the role of this molecule is likely to be dependent on aggregate clearance and overall effects on proteostasis. Using this platform we identified harman, a naturally occurring β-carboline alkaloid present in cooked foods and tobacco smoke, as a reproducible reducer of PFF-induced aggregation. Conversely, compounds including amsacrine and carnosol were identified as aggregation promoting compounds. These findings support the hypothesis that small molecules derived from environmental or microbial sources can directly influence aggregation dynamics in live cells.

Environmental exposures and dietary components have been widely linked to PD risk. Interestingly, compounds such as nicotine and caffeine have been associated with reduced PD risk ^33,34^, and non-caffeine and non-nicotine phenolic compounds found in coffee and cigarette smoke have been reported to have neuroprotective effects ^35^. Our identification of harman as an inhibitor of aggregate accumulation may represent a mechanistic example of protective roles for certain bioactive dietary molecules. At the molecular level, our transcriptomic studies showed that changes in thioredoxin peroxidase activity were associated with harman treatment. It is interesting to note that periredoxins have been associated with neuronal death in PD ^36,37^ but also with inflammation and lysosomal disruption in the intestine ^38^. In our assay, knockdown of PRDX5 alone, a unique peroxiredoxin with broad subcellular localization ^39^, was sufficient to recapitulate the effect of harman of synuclein aggregate formation within HEK293T cells. While periredoxins are broadly expressed, it is important to note that HEK293T cells, while amenable to reproducible high-throughput screening, may not properly model neuronal synuclein aggregation or its effect on dopaminergic neuron biology, which is likely subset specific ^40^. Neuronal models would provide greater physiological relevance but are less amenable to high-throughput imaging pipelines due to their morphology and variability. We note however that α-syn is expressed in both neuronal and non-neuronal contexts, and increasing evidence supports roles for peripheral cell types in disease initiation and propagation ^41–45^. In particular, gut associated cells such as enteroendocrine cells, have been implicated in α-syn transmission and early pathology, suggesting that non-neuronal systems can capture biologically relevant aspects of aggregation, particularly in body-first models of PD ^41,45^, where peripheral nervous system involvement precedes central motor symptoms ^10,11,46^.

At the chemical level, our study focused primarily on metabolites with validated standards detected in the Human Microbiome Project, but many emerging studies have focused on expanding the annotated chemical space and linking specific microbial taxa to the metabolic products they generate ^23,47^. Extending these assays to metabolites associated with PD-enriched or PD-depleted microbes is likely to reveal novel actionable modulators of synuclein aggregation and to better define environmental factors involved in the onset and progression of PD.

## Methods

### Cell Culture

HEK293T cells (ATCC Cat# CRL-3216, RRID:CVCL_0063) were grown in sterile-filtered DMEM (Thermo Fisher Scientific, 10569044) supplemented with 10% heat-inactivated FBS (ATCC, 30-2020), 10 mM HEPES (Thermo Fisher Scientific, 15630080), and 10,000 U/mL penicillin-streptomycin (Thermo Fisher Scientific, 15140122). Cells were incubated at 37°C, 5% CO_2_.

### Generation of Stable HEK293T Cell Line expressing GFP-α-synuclein

HEK293T cells were seeded at 3 x 10^5^ cells per well in 3 mL of supplemented DMEM in sterile 6-well tissue culture-treated plates and incubated overnight. EGFP-alphasynuclein-WT plasmid (gift from David Rubinsztein, Addgene #40822) was transfected with lipofectamine according to the manufacturer’s guidelines (Thermo Fisher Scientific, L3000001).

Cells were incubated for 48h, then medium was supplemented with G418 Sulfate (Geneticin, Thermo Fisher Scientific, 10131035) selective antibiotic at 500 µg/mL. Cells were incubated for two weeks to select for stable expressors, then sorted by GFP expression on a FACS-based cell sorter (Sony Biotechnology).

Cells in the highest-expressing 0.72% of the population were plated into individual wells of sterile 96-well TC-treated flat-bottom plates in 150 µL supplemented DMEM. 24h after sorting, G418 was added at 500 µg/mL. Cells were expanded from individual clones for three weeks and GFP expression was validated, after which working stocks of selected clones were frozen in 10% DMSO in liquid nitrogen. Data analysis was performed in FlowJo (Version 10.10.0; https://www.flowjo.com/flowjo10/overview, RRID:SCR_008520). A-syn-GFP-mCherry line was generated following the same procedure as outlined above by cloning in mCherry/Puro plasmid (Vector Builder VB010000-9290tfx).

### Transfection of Pre-Formed α-synuclein Fibrils (PFFs) and Aggregation-Modulating Compounds into Cell Cultures

The protocol used for treatment and imaging is available with the following accession number: https://dx.doi.org/10.17504/protocols.io.5jyl8p7wrg2w/v1

24h prior to transfection, α-syn-GFP HEK293T cells were seeded in replicate 384-well poly-D-lysine coated optical plates (Revvity, 6057500) at a density of 5000-8000 cells per well in 40 µL culture medium (without G418) and incubated overnight.

Prior to transfection, Lipofectamine 3000 was added to Opti-MEM medium at a 5% (v/v) concentration and incubated at room temperature for 5 minutes. PFFs (StressMarq, SPR-322) were thawed on ice for 15 minutes. Thawed PFFs were diluted in Opti-MEM medium and sonicated in Covaris milliTUBEs (Covaris, 520128) at room temperature using a Covaris LE220Rsc focused ultrasonicator (2 bursts, 65 seconds / burst, 1000 cycles / burst, 75W peak power, 15W average power, 20% duty factor) ^48,49^. Sonicated PFFs were diluted to 40 µg/mL in Opti-MEM medium, then mixed with Lipofectamine 3000/Opti-MEM solution and incubated at room temperature for 20 minutes.

For transfection, 10 µL of this PFF-Lipofectamine mixture was added to each well, resulting in a final well volume of 50 µL and a final PFF concentration of 4 µg/mL. Negative control wells received 10 µL of 2.5% Lipofectamine 3000 in Opti-MEM without PFFs. Immediately following transfection, plates were centrifuged at 100 x g for 4 minutes at room temperature.

400 nL of each compound (1-10 mM stock concentration in DMSO, Sigma-Aldrich, D5879-100ML) was added after transfection using a robotic pin tool liquid handling system via four pinning cycles using a 100 nL pin tool attachment, with three wash cycles between each pin (AnalytikJena CyBio Pintool). Negative control wells received DMSO only. All plates were kept at 37°C during the procedure with the exception of the plate currently being pinned at room temperature. Upon completion, plates were returned to the incubator for 24h prior to fixation and staining.

### Fixation and Nuclear Staining

Cells were fixed by adding EM-grade paraformaldehyde (Electron Microscopy Sciences, 15714) to a final concentration of 4%. Plates were centrifuged at 100 x g for 5 minutes at room temperature, then incubated at room temperature for 15 minutes. Medium was aspirated, and using a plate washer and liquid handling control (Agilent BioTek EL406) 20 µL of 1x DPBS containing 8 μg/mL Hoechst 33342 nuclear stain (Invitrogen, H3570) was added directly to cells. Cells were incubated at room temperature for 15 minutes, washed once with 1x DPBS (Sigma-Aldrich D8537), and stored in 50 µL of DPBS at 4°C protected from light until imaging.

### High-Throughput Cell Imaging

Cells were imaged directly in 384-well plates using the Opera Phenix Plus High-Content Screening System (Revvity). Nine images per well were captured in three planes (2 μm Z-stack) using a 20x confocal water objective lens. α-syn-GFP signal was captured using the green fluorescence channel (major emission wavelength: 509 nm). Hoechst 33342 signal was captured in the blue fluorescence channel (major emission wavelength: 461 nm). When working with the α-syn-GFP-mCherry cell line, the mCherry signal was captured with the red fluorescence channel (major emission wavelength: 610 nm).

### Filtering and Quantification of Intracellular Puncta

A custom analysis pipeline was developed in Harmony software (Revvity, Version 5.2, https://www.perkinelmer.com/uk/product/harmony-4-8-office-hh17000001, RRID:SCR_018809) and implemented in SImA (Version 1.4.0.29; https://revvitysignals.com/image-artist, RRID:SCR_024466) to identify cells and quantify intracellular α-syn aggregates. Nuclei were first detected using Hoechst 33342 to identify and count individual cells. Cytoplasmic boundaries were defined based on GFP fluorescence, with cells at the image edges excluded from analysis. Mean per-cell GFP positivity was calculated, and low-expressing cells below a predefined threshold were removed.

GFP-positive puncta within the cytoplasmic boundaries of the remaining cells, representing possible α-synuclein aggregates, were filtered by size, contrast, and background intensity, and debris erroneously called as cells was excluded. The remaining puncta were considered to be “true” α-synuclein aggregates. The effects of modulating compounds were quantified by calculating the per-cell number and area of these puncta, averaged over each well. Statistical analyses on the extracted data were performed in Prism (GraphPad, Version 10.6.1, www.graphpad.com/scientific-software/prism/, RRID:SCR_002798) as detailed in each figure legend.

### Treatment of HEK293T Cells with Harman for RNA Isolation

Fresh HEK293T and α-syn-GFP-mCherry stocks were thawed and grown to confluence. Cells were harvested and seeded into 12-well plates at 4 x 10^5^ cells per well and incubated overnight to reach a confluency of 80-90%. Cells were transfected with PFFs and compounds were added as described above, with volumes scaled to a 1mL final volume for a 12-well plate format. Cells were returned to 37°C, 5% CO_2_ for 24h.

### RNA Extraction for Bulk RNA-Sequencing

RNA was extracted with TRIzol reagent as per manufacturer’s instructions (Invitrogen 15596062). The quality of isolated RNA samples was assessed using Agilent HS RNA ScreenTape (Agilent, 5067-5579) on the Agilent 4200 TapeStation.

### SmartSeq2 based Bulk RNA sequencing

To obtain full-length transcriptomic data from cultured cells we performed SmartSeq2-based bulk RNA sequencing as in ^50^. In brief, RNA lysate clean-up was performed with Agencourt RNAClean XP Beads (Beckman Coulter, A639867). Next, Maxima H minus reverse Transcriptase (Thermo Fisher Scientific, EP0753) was used for reverse transcription and KAPA HotStart HIFI PCR ReadyMix (KAPA Biosystems, KK2602) was used for whole-transcription amplification (WTA). WTA products were purified with Ampure XP DNA SPRI beads (Beckman Coulter, A63881) and quantified using Qubit dsDNA HS Assay Kit (Thermo Fisher Scientific, Q32851). Quality was assessed with D5000 HS ScreenTape (Agilent, 5067-5588) on an Agilent 2200 TapeStation. WTA products were diluted for tagmention and indexing with the Nextera XT DNA Library Preparation Kit (Illumina, FC-131-1096) and two subsequent clean-ups with Ampure XP DNA SPRI beads. For quality control, Qubit dsDNA HS Assay Kit and Agilent D1000 ScreenTape (Agilent, 5067-5584) were used to determine concentration and size. Samples were pooled together and sequenced on a NextSeq 2000 (Illumina) using a P2 flow cell and the following read configuration: R1, R2 = 38bp, I1, I2 = 8bp.

### SmartSeq2 FASTQ Processing

The flow cell data was demultiplexed with bcl-convert (Illumina, Version 4.2.7, https://support.illumina.com/downloads/bcl-convert-v4-2-7-installers.html) and quality control was performed using FastQC (Babraham Bioinformatics, Version 0.11.5, http://www.bioinformatics.babraham.ac.uk/projects/fastqc/, RRID:SCR_014583). Paired FASTQs were pseudoaligned and counted using kallisto ^51^ (Version 0.46.1, https://pachterlab.github.io/kallisto/about, RRID:SCR_016582) and a 31-mer index built from the reference human transcriptome GRCh38. Resulting h5 files were imported into R (Version 4.3.2, https://www.r-project.org/, RRID:SCR_001905) using the Tximport ^52^ package (Version 1.38.2, https://github.com/mikelove/tximport, RRID:SCR_016752) and counts values were normalized and aggregated at the gene level during import.

### Transcriptomic Analysis

Differential expression analysis was performed using DESeq2 ^53^ (Version 1.42.1, http://bioconductor.org/packages/release/bioc/html/DESeq.html, RRID:SCR_000154). Counts were modeled using the following design formula:

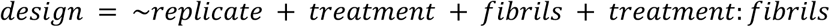

Differential expression for each term was assessed using Wald tests. For pairwise contrasts, results were analyzed in the absence vs. presence of harman, the absence vs. presence of PFFs, and the treatment:fibril interaction, capturing genes whose response to harman is altered in the presence of PFFs. Genes with FDR < 0.2 were considered significant.

For each term, the Wald statistic calculated for all genes was used to assess correlations of transcriptional responses between cell lines (**Supplementary Table 3)**. Pearson correlation coefficients were calculated, and a linear fit was overlaid for visualization.

### Enrichment Analysis

Differentially expressed gene (DEG) sets (**Supplementary Table 3**, padj < 0.2) were uploaded to the PAN-GO Human Functionome Molecular Function tool (PAN-GO, Version 2.0, https://functionome.geneontology.org, Released 2025-10-12) and pathway enrichment analysis was performed using Fisher’s Exact test with Benjamini-Hochberg correction (FDR ≤ 0.05). Up-and downregulated pathways with a fold enrichment (number of genes detected / number of genes expected based on group size) ≥ 5 were included to prioritize robust signals.

### Aggregation Screens into siRNA Treated Cell Cultures

96-well PhenoPlates (Revvity, 6055302) were coated with poly-lysine solution (Poly-L-lysine, Sigma Aldrich, P8920, used at 0.01% w/v or Poly-D-Lysine, Gibco, A38904-01, 0.1 mg/mL) prior to seeding. 50μL of poly-lysine solution was added to each well and plates were incubated for 10 minutes at 37°C, 5% CO2, then washed 4x with H2O and dried for at least 4h in the biosafety cabinet. Plates were wrapped in parafilm and stored at 4°C until use.

siRNAs (**Supplementary Table 1**) were reconstituted in nuclease-free water to 100 μM and diluted to 10 μM in nuclease-free duplex buffer (IDT, 11-01-03-01). Prior to transfection, siRNA was further diluted to 1 μM working stock for transfection in nuclease-free water.

siRNAs were reverse transfected into the α-syn-GFP-mCherry cell line with Lipofectamine RNAiMAX Transfection Reagent (Thermo Fisher, 13778030) per manufacturer instructions, scaled to a final volume of 160 μL per well and a final siRNA concentration of 10 nM.

Individual siRNA conditions included negative scrambled control (NC), PRDX1.1, PRDX1.2, PRDX2.1, PRDX2.2, PRDX5.1, PRDX5.2, and PRDX5.3. For each transfection condition, siRNA-lipid master mixes were prepared in bulk to dispense into replicate wells. In brief, 1.6 pmol of siRNA was diluted in 25.13 μL of Opti-MEM and combined with 0.27 μL Lipofectamine RNAiMAX per well. Complexes were incubated at room temperature for 20 minutes and added to culture wells.

α-syn-GFP-mCherry cells were diluted in complete growth medium without antibiotics where 133 μL contained 9,000-12,000 cells and was added to each well containing siRNA-lipid complexes. This resulted in a final volume of 160 μL and final RNA concentration of 10 nM for each individual siRNA. To mix, the plate was gently rocked back and forth and incubated at 37°C, 5% CO_2_.

Two plates per screen were prepared: one for the optical screen and one for validation experiments. For the optical screen, six technical replicate wells were prepared for individual conditions, in addition to no-siRNA control wells. A separate validation plate with the same individual siRNA conditions was prepared in triplicate.

2 days post transfection, cells were transfected with PFF as described previously at a final concentration of 2 μg/mL. Harman was added in triplicate to individual and combinatorial siRNA wells and to no-siRNA control wells at a final concentration of 80 μM and incubated at 37°C, 5% CO_2_.

24h later, 72h following transfection, cells are fixed and stained as described prior scaled to a 96-well format.

Cells were imaged as described above (*High-Throughput Cell Imaging* section) in 96-well plates using the Opera phenix Plus High-Content Screening System, the only change being 25 images per well were captured vs. 9 images per well in 384-well plates.

### Validation of siRNA Activity with RT-qPCR

2 days after transfection, RNA was extracted from the “validation” plate with the Quick-RNA 96 kit (Zymo, R1052) in an RNase free AirClean Systems PCR Workstation. RNA concentration was measured with the Qubit RNA High Sensitivity (HS) Assay Hit (Thermo Fisher, Q32855).

For quantitative PCR, 25 ng of RNA and respective primer pairs (**Supplementary Table 1**) were added to iTaq™ Universal SYBR® Green One-Step master mix (Bio-Rad, 1725150).

Measurements were conducted with the CFX96 Real-Time PCR Detection System (Bio-Rad).

Fold-change change between siRNA treated and negative scrambled controls were calculated as 2^−ΔΔ*C_t_*^ using *HPRT* as a house-keeping gene and expressed as a percentage of knockdown.

## Acknowledgments and author contributions

This research was funded in part by Aligning Science Across Parkinson’s [ASAP-000529] through the Michael J. Fox Foundation for Parkinson’s Research (MJFF). For the purpose of open access, the author has applied a CC BY public copyright license to all Author Accepted Manuscripts arising from this submission. We thank Maria Alimova and the Center for the Development of Therapeutics (CDoT) at the Broad Institute for assistance with the establishment of the imaging and analysis pipeline.

L.R., C.G., A.B. and J.L. performed experiments and analyzed data under supervision from K.C., J.D. and R.J.X; L.R. and J.D. wrote the initial draft of the manuscript; K.C., J.D. and R.J.X conceptualized the work; all authors reviewed and edited the final version of the manuscript.

R.J.X. is co-founder of Jnana Therapeutics and Convergence Bio, Board Director at MoonLake Immunotherapeutics, consultant to Nestlé, and a member of Magnet Biomedicine and Arena Bioworks’ scientific advisory boards; J.D. is a member of Biorender’s scientific advisory board; these organizations had no role in this study. All other authors declare no competing interests.

## Data availability

The raw and processed transcriptomic data is publicly available on GEO, accession number GSE328400.

---

**Supplementary Figure 1, related to Figure 1.**
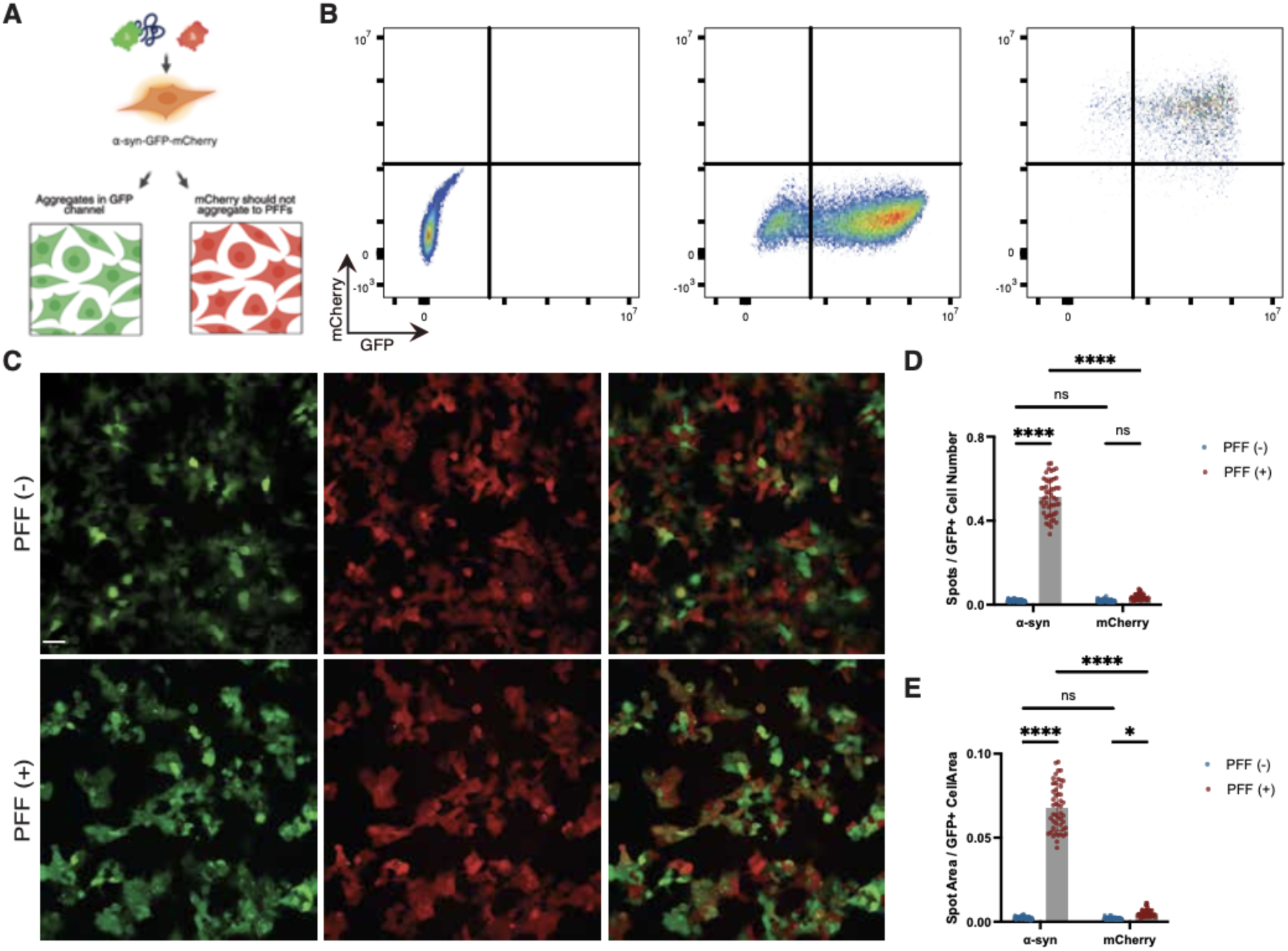
Validation of aggregation specificity using an α-syn-GFP-mCherry reporter Cell Line. **(A)** Schematic of α-syn-GFP-mCherry cell line construct created with Biorender.com. **(B)** Flow cytometry analysis used to generate a stable α-syn-GFP-mCherry HEK293T cell line. Shown are representative FACS plots from GFP-negative wild type (WT) controls (left), α-syn-GFP positive cells (middle), and α-syn-GFP-mCherry positive cells (right). **(C)** Representative confocal images of untreated (top) and PFF-treated (bottom) α-syn-GFP-mCherry cells. GFP puncta (purple) and mCherry puncta (blue) are labeled. Scale bar, 50 *μ*m. **(D-E)** Quantification of puncta formation in GFP and mCherry channels. Comparison of α-synuclein puncta and mCherry puncta shown as puncta per cell (D) and puncta area normalized to cell area (E). Data were analyzed by one-way ANOVA with Dunnett’s T3 multiple comparisons post-hoc test.; ****p < 0.0001; n = 48 wells per group; pooled from 4 independent experiments.

**Supplementary Figure 2, related to Figure 2.**
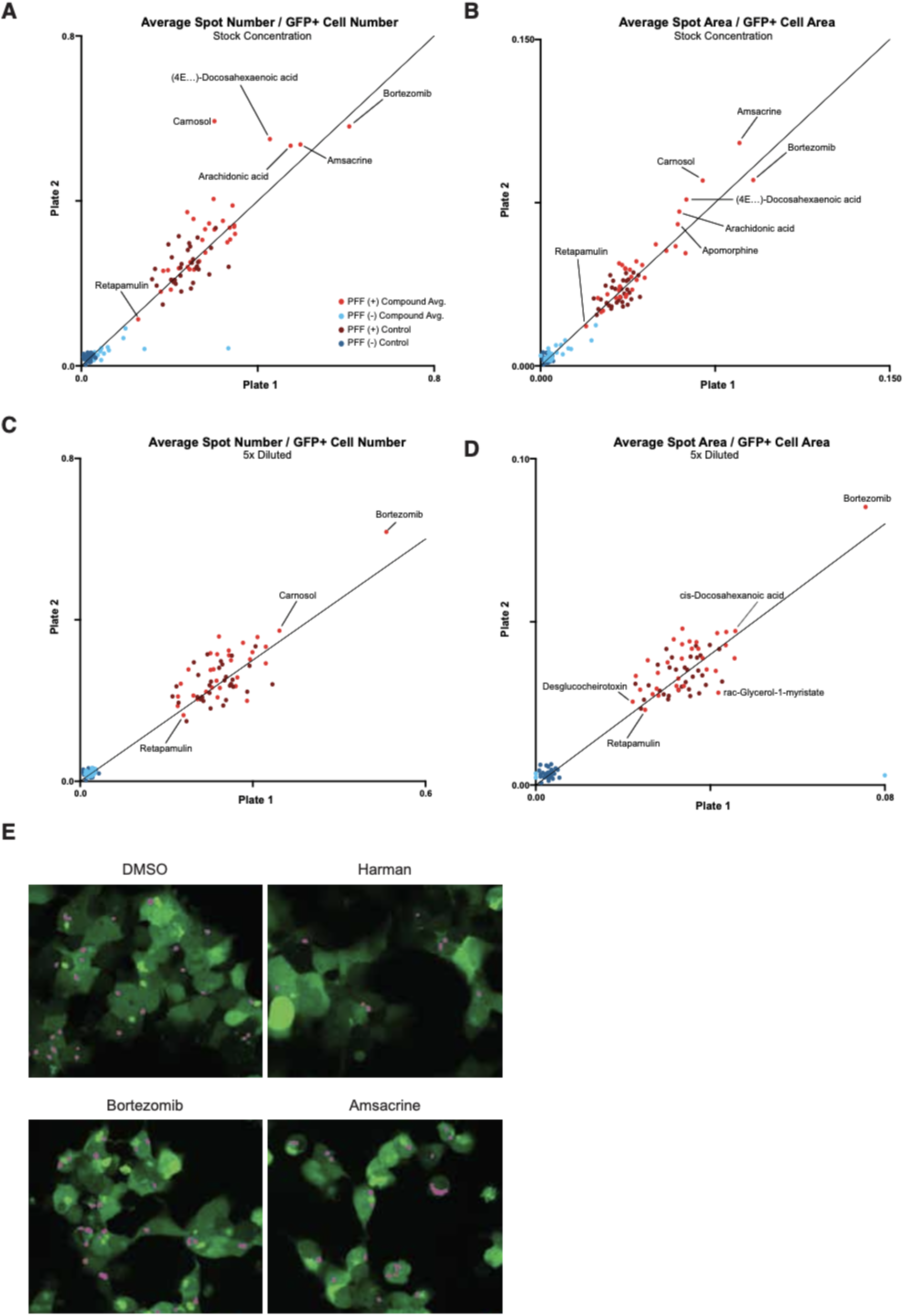
Dose response for selected compounds. Quantification of the impact on spot formation of 32 microbiome and gut-related compounds at stock concentration (8-80 μM) **(A-B)** and 5x diluted concentration (1.6-16 μM) **(C-D)** across replicate plates; intracellular GFP puncta are quantified as GFP spot number normalized to cell number **(A,C)** and GFP spot area normalized to cell area **(B,D)**. Each data point represents the average of 6 wells. **(E)** Representative confocal images microscopy images of select compounds shown at higher magnification from the corresponding panels in Fig. 2F.

**Supplementary Figure 3, related to Figure 3.**
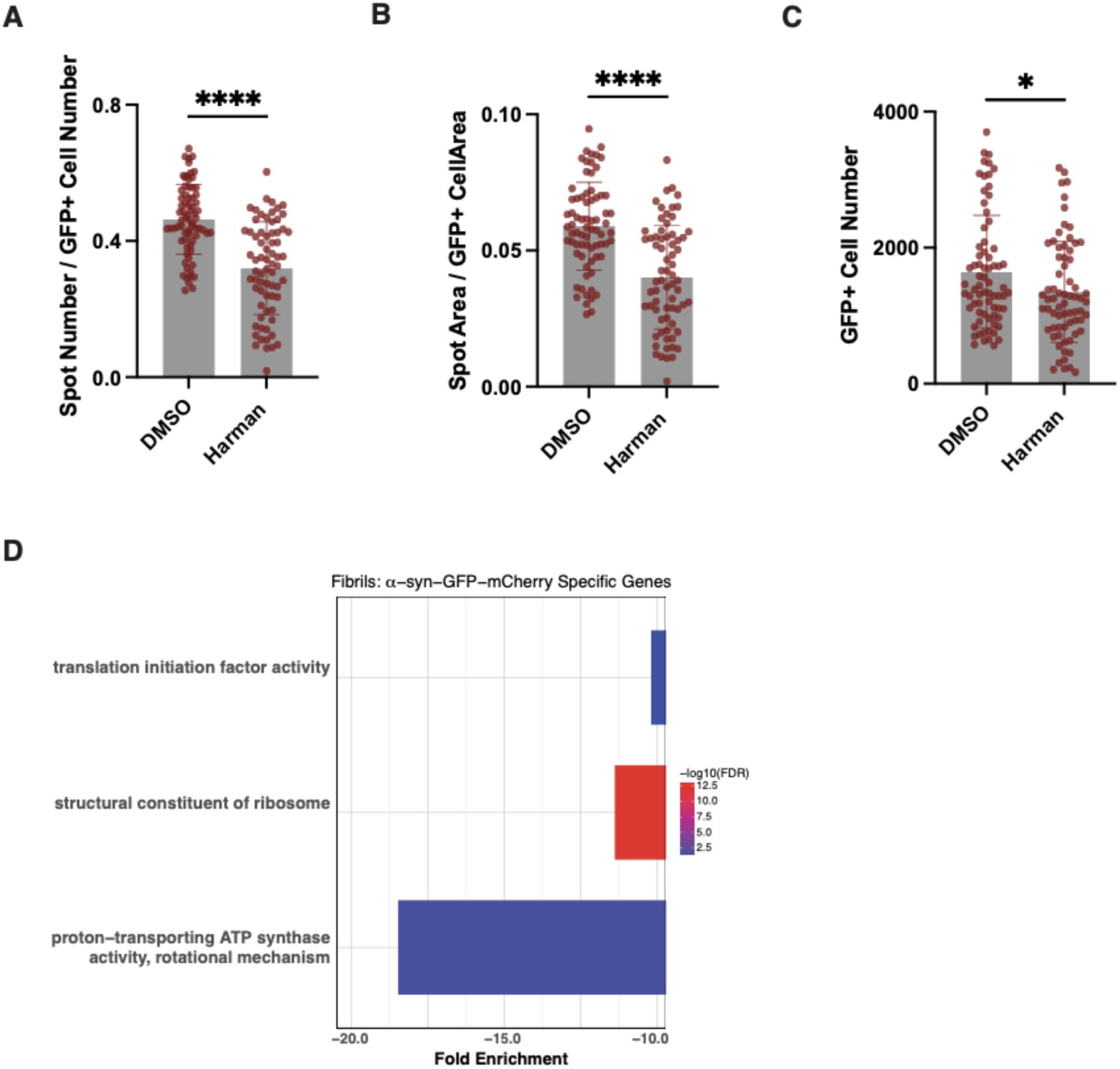
Transcriptional impact of PFF treatment on synuclein expressing and control cells (A-C) Quantification of GFP puncta number per cell **(A)**, GFP puncta area per cell area **(B)**, and GFP cell number **(C)** in cells treated with PFFs and DMSO (n = 72 wells) or PFFs and Harman (n = 69 wells). Unpaired two-tailed t-test with Welch’s correction: *p < 0.05 **p<0.01, ****p < 0.0001; 4 independent experiments. **(D)** Functionome GO molecular function tool pathway of specific genes significantly downregulated in α-syn-GFP-mCherry cells (FDR < 0.2 and fold change < 0) but not in control cells (untested or FDR ≥ 0.2)

## Other supplementary materials

Supplementary Table 1, related to Methods. Key resources table including reagents, software, oligonucleotide sequences and datasets

Supplementary Table 2, related to Figure 2. Chemical compound identity and screening results

Supplementary Table 3, related to Figure 3. RNA sequencing results For each term (effect of harman, fibrils or interaction term) in α-syn-GFP-mCherry or wild-type cells, results of DESeq2 analysis are provided (gene - gene symbol; baseMean - base mean expression; log2FoldChange - log2 fold change estimate associated with the indicated term; lfcSE - log2 fold change standard error estimate; stat - Wald statistic; pvalue - nominal p value; padj - corrected p value)

